# Divergent microbiota of echinoid eggs separated by the Isthmus of Panama

**DOI:** 10.1101/2020.03.30.015578

**Authors:** Tyler J. Carrier, Harilaos A. Lessios, Adam M. Reitzel

## Abstract

Relationships between animals and their associated microbiota is dependent on the evolutionary history of the host and on the environment. The majority of studies tend to focus on one of these factors and rarely consider how both determine the community composition of the associated bacteria. One “natural experiment” to test how evolutionary history, shared environments, and the interaction between these factors drive community composition is to compare geminate species pairs. Echinoids separated by the Isthmus of Panama are suitable for this comparison due to the known evolutionary history and differences in oceanographic characteristics of the Caribbean Sea and Pacific Ocean. By comparing the egg-associated microbiota for the *Echinometra* and *Diadema* geminate species pairs, we show that both pairs of geminate species associate with distinct bacterial communities in patterns consistent with phylosymbiosis, and that the interaction between the evolutionary history of the host and the environment best explain differences in these communities. Moreover, we find that particular microbial taxa differed considerably between, but not within, oceans and that the microbiota of the two Caribbean *Echinometra* species were dominated by the phototrophic Oxyphotobacteria.

## INTRODUCTION

Relationships between animals and the microbes with which they associate is generally hypothesized to be driven by two primary factors, the evolutionary history of the host and the environment (Bordenstein & Theis 2015). In the former, animals associate with species-specific microbial communities that often co-vary with host phylogeny (Schmitt et al. 2012, Brooks et al. 2016, Carrier & Reitzel 2018, Pollock et al. 2018, O’Brien et al. 2019, Lim & Bordenstein 2020). In the latter, the composition of host-associated microbiota depends on the abiotic environment, such that bacterial communities may differ due to bacterial physiology or characteristics of the host to form a community with specific functional properties (Soto et al. 2009, Webster et al. 2011, Kohl & Carey 2016, Carrier & Reitzel 2017).

Studies assessing the relative importance of host’s evolutionary history or the environment suggest that both influence community composition, but one factor is commonly more pronounced. In scleractinian corals (Pollock et al. 2018) and sponges (Easson & Thacker 2014, Thomas et al. 2016), for example, host phylogeny can best explain compositional differences in these microbial communities. Shifts in symbiont composition due to the abiotic environment have been observed for some echinoderms (Carrier & Reitzel 2020), sponges (Webster et al. 2011), and barnacles (Aldred & Nelson 2019). In select cases, however, the influence of host’s evolutionary history and the environment are comparably similar (Mortzfeld et al. 2015).

Discerning the impacts of evolutionary history and adaptation to different environmental conditions is a challenge in many marine species because of incomplete or unknown phylogenetic relationships and overlapping environments for closely related species. Geminate species, or sister species pairs resulting from a recent speciation event following a geological event (Jordan 1908), represent insightful systems to compare between these factors to determine how they relate to the associated microbiota (Wilkins et al. 2019). One geographic change that resulted in multiple geminate species was the formation of the Isthmus of Panama. Until ~2.8 million years ago (MYA), the Caribbean Sea and Tropical Eastern Pacific was a continuous water basin with fauna that spanned this region (Lessios 2008, O’Dea et al. 2016). The emergence of the Isthmus of Panama affected the physical conditions of these two oceans, causing multiple groups of marine fauna to undergo independent evolutionary trajectories and to form geminate species pairs (Lessios 2008, O’Dea et al. 2016, Wilkins et al. 2019).

Two sets of geminate species that have been particularly well-studied are the echinoids *Echinometra* and *Diadema*. As a result of the formation of the Isthmus of Panama, *Diadema* split into the Pacific *D. mexicanum* and Caribbean *D. antillarum* (Lessios et al. 2001, Hickerson et al. 2006, Lessios 2008). *Echinometra* also diverged into *E. vanbrunti* in the Pacific and *E. lucunter* and *E. viridis* in the Caribbean, with the speciation event for these two Caribbean species occurring after the emergence of the Isthmus of Panama (McCartney et al. 2000, Lessios 2008). Fundamental differences in egg size and biochemical composition (Lessios 1990, McAlister & Moran 2012), larval feeding ecology (McAlister 2008), and reproductive ecology (Lessios 1981, 1984) have occurred since these geminate pairs diverged.

The microbiota associated with geminate species pairs was recently hypothesized to have diverged during their independent evolutionary trajectories, either as a product of the host genetics or environmental differences (Wilkins et al. 2019). For the echinoid geminate species pairs, relatedness of microbial communities would be hypothesized to either mirror host phylogeny or form distinct clades unique to each ocean, regardless of phylogeny (Fig. 1). To test these hypotheses, we sampled eggs of all five *Echinometra* and *Diadema* species and used amplicon sequencing to profile the associated bacterial communities.

**Fig. 1:**
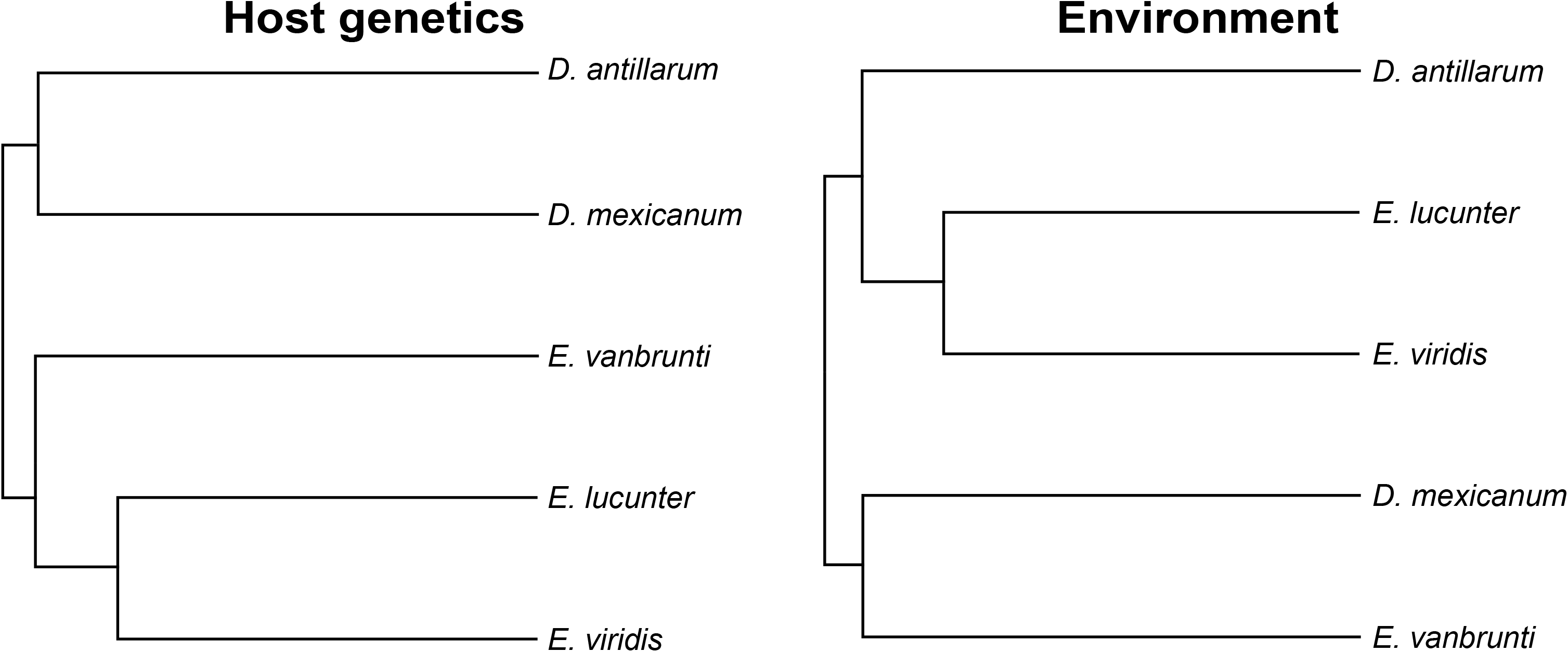
Hypothesized relationships of microbial communities for geminate species pairs in this study. Hypothetical microbial dendrograms for *Echinometra* and *Diadema* geminate species pairs if host genetics (left) or the environment (right) were the primary factor contributing to community assembly.

## MATERIALS AND METHODS

### Echinoid collections and spawning

Adult *Diadema mexicanum* and *Echinometra vanbrunti* were collected by SCUBA off Isla Taboguilla near Panama City, Panama. Adults were transferred to and spawned at the Smithsonian Tropical Research Institute’s (STRI) Naos Island Laboratories in Balboa. *D. antillarum*, *E. lucunter*, and *E. viridis* were collected at Galeta Marine Laboratory near Colón, Panama in the Caribbean Sea. Of the Caribbean species, *E. lucunter* and *E. viridis* were brought to and spawned at Naos, while *D. antillarum* was spawned at Galeta.

Adult sea urchins (n=15 per species; Table S1) were spawned by intracoelomic injection of 0.50 M KCl. For all three *Echinometra* species, eggs were shed into 5.0 μm filtered seawater from their site of origin while *Diadema* eggs for both species were shed onto and collected from the aboral surface. Approximately ~250 eggs per individual were collected with a sterile pipette, concentrated using a microcentrifuge, preserved in RNAlater, and frozen at −20°C for long-term storage.

The environmental microbiota from the seawater where the sea urchins were collected was also sampled. For both Pacific and Caribbean populations, ~0.5-L of seawater was filtered onto a 0.22-μm Millipore filter to retain the environmental microbiota (n=3). Full filter disks were preserved in RNAlater and frozen at −20°C.

### Profiling bacterial communities

Total DNA was extracted from sea urchin eggs, seawater samples, and DNA kit/ reagent blanks (n=3) using the GeneJet Genomic DNA Purification Kit (Thermo Scientific). DNA was quantified using a Qubit Fluorimeter (Life Technologies) and diluted to 5 ng•μL_−1_ using RNase/DNase-free water. Bacterial sequences were then amplified using primers for the V3/V4 regions of the 16S rRNA gene (Table S2; Klindworth et al. 2013). Products were purified using the Axygen AxyPrep Mag PCR Clean-up Kit (Axygen Scientific), indexed using the Nextera XT Index Kit V2 (Illumina Inc.), and then purified again. At each clean up step, flxuorometric quantitation was performed using a Qubit, and libraries were validated using a Bioanalyzer High Sensitivity DNA chip (Agilent Technologies). Illumina MiSeq sequencing (v3, 2×300 bp paired-end reads) was performed in the Department of Bioinformatics and Genomics at the University of North Carolina at Charlotte.

### Computational analysis

Raw reads along with quality information were imported into QIIME 2 (v. 2019.1; Bolyen et al. 2019), where adapters were removed, forward and reverse reads were paired using VSEARCH (Rognes et al. 2016), filtered by quality score, and denoised using Deblur (Amir et al. 2017). QIIME 2-generated ‘features’ were analyzed as Amplicon Sequence Variants (ASVs; Callahan et al. 2017) and were assigned taxonomy using SILVA (v. 132; Quast et al. 2013). Sequences matching to Archaea, present in the seawater or in DNA kit/ reagent blanks were filtered from the data table, and samples with less than 1,000 reads were discarded (Table S1). The filtered table was then rarified to 1,027 sequences per sample (*i.e.*, the read count for the sample with the least remaining reads; Fig. S1),

To test whether community membership and composition were species-specific, we calculated unweighted and weighted UniFrac values (Lozupone & Knight 2005) and compared them using principal coordinate analyses. Results from these analyses were then recreated in QIIME 1 (v. 1.9.1; Caporaso et al. 2010) and stylized using Adobe Illustrator. We then used a PERMANOVA to test for differences in membership and composition between species and, subsequently, performed pairwise comparisons. We also calculated several measures of alpha diversity: total ASVs, Faith’s phylogenetic diversity, McIntosh dominance, and McIntosh evenness. We compared these values using a one-way analysis of variance (ANOVA) for *Echinometra* and a Student’s t-test for *Diadema*. Lastly, we summarized the bacterial groups associated with all species and the number of shared and species-specific ASVs.

Using the weighted UniFrac values, we constructed microbial dendrograms in QIIME 2 and tested for phylosymbiosis by comparing topological congruency with a portion of the cytochrome c oxidase subunit 1 (CO1) gene tree for these geminate species pairs (McCartney et al. 2000, Lessios et al. 2001). The host CO1 tree was constructed using BEAST (v. 1.8.4; Drummond et al. 2012) by starting from a random coalescent tree and running for 10_7_ steps, with recordings every 10_3_ steps. We then used Tracer (v. 1.7.1; Rambaut et al. 2018) to verify that Effective Sample Size values for all parameters exceeded 4,260. A maximum credibility tree, with support of 1 for all nodes, was then estimated using TreeAnnotator (v. 10.10.4). Patterns of phylosymbiosis were tested using the Robinson-Foulds metric in TreeCmp (Bogdanowicz et al. 2012) and matching cluster metrics with 10,000 random trees. This analysis was performed using the Python script of Brooks et al. (2016).

Our QIIME-based pipeline used to convert raw reads to ASVs for visualization is presented in detail in Supplemental Note 1. The 16S rRNA gene reads are accessible on Dryad Digital Repository (doi.org/10.5061/dryad.2z34tmphq).

## RESULTS

### Community relatedness and diversity

The bacterial communities associated with eggs of these two geminate species pairs were species-specific in both community membership and composition, except in one case (unweighted UniFrac: p<0.001; weighted UniFrac: p<0.001; Fig. 2; Table S3). The two Caribbean *Echinometra* species, *E. lucunter* and *E. viridis*, associated with comparatively similar communities in both membership and composition (unweighted UniFrac p=0.05; weighted UniFrac: p=0.095; Fig. 2; Table S3). Relatedness of these bacterial communities suggests that the Pacific *E. vanbrunti* groups with the Pacific *D. mexicanum* and Caribbean *D. antillarum* while the Caribbean *E. lucunter* and *E. viridis* group separately (Fig. 2). Moreover, comparison of host phylogeny and bacterial dendrogram suggest that these are congruent and, thus, phylosymbiosis was supported for these geminate species pairs (Robinson-Foulds: p=0.062; Fig. 3; Table S4).

**Fig. 2:**
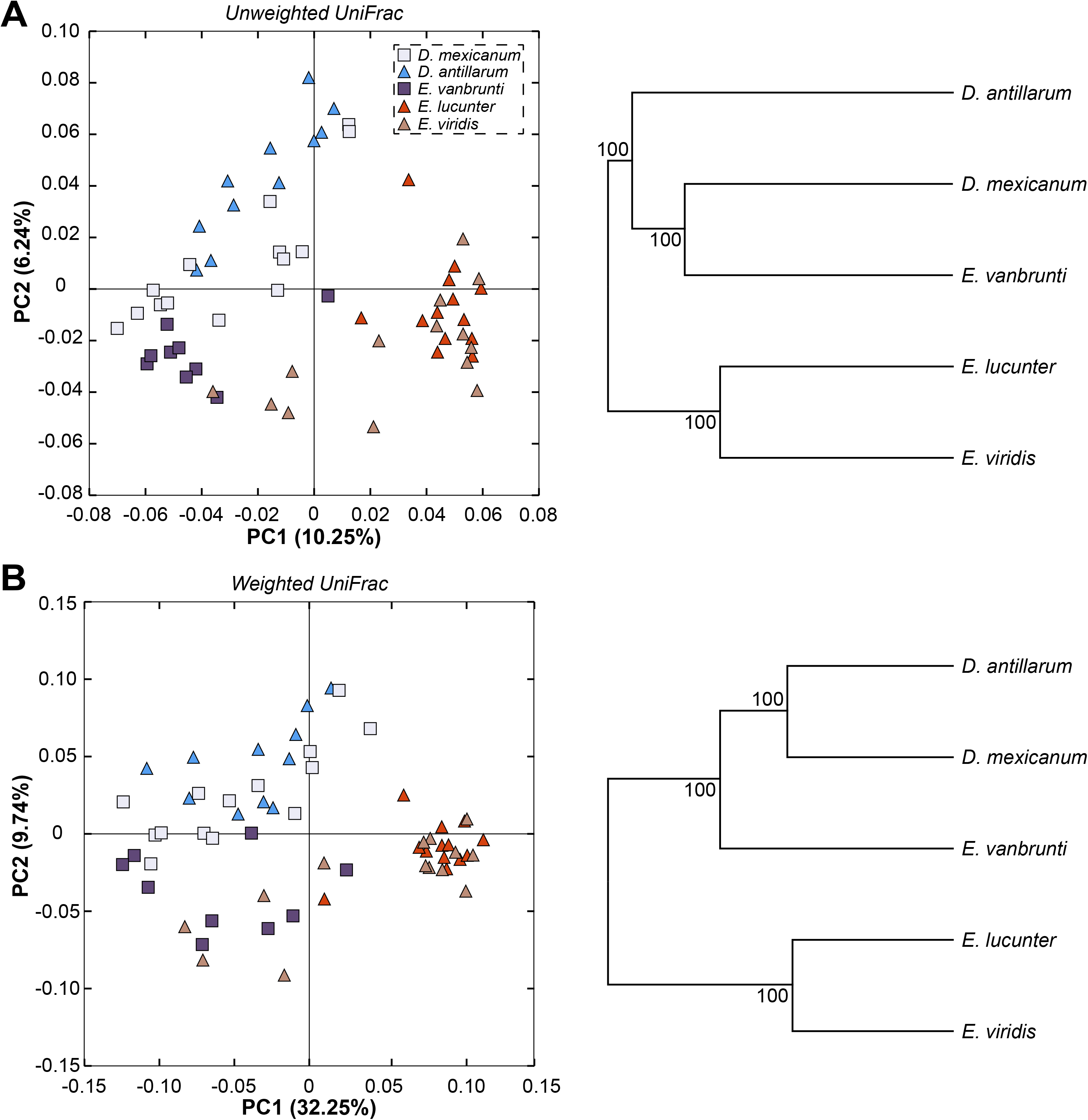
Community relatedness of geminate species pairs. Similarity between the bacterial communities of *Echinometra* and *Diadema* geminate species pairs based on membership (unweighted UniFrac; A) and composition (weighted UniFrac; B) with corresponding microbial dendrograms.

**Fig. 3:**
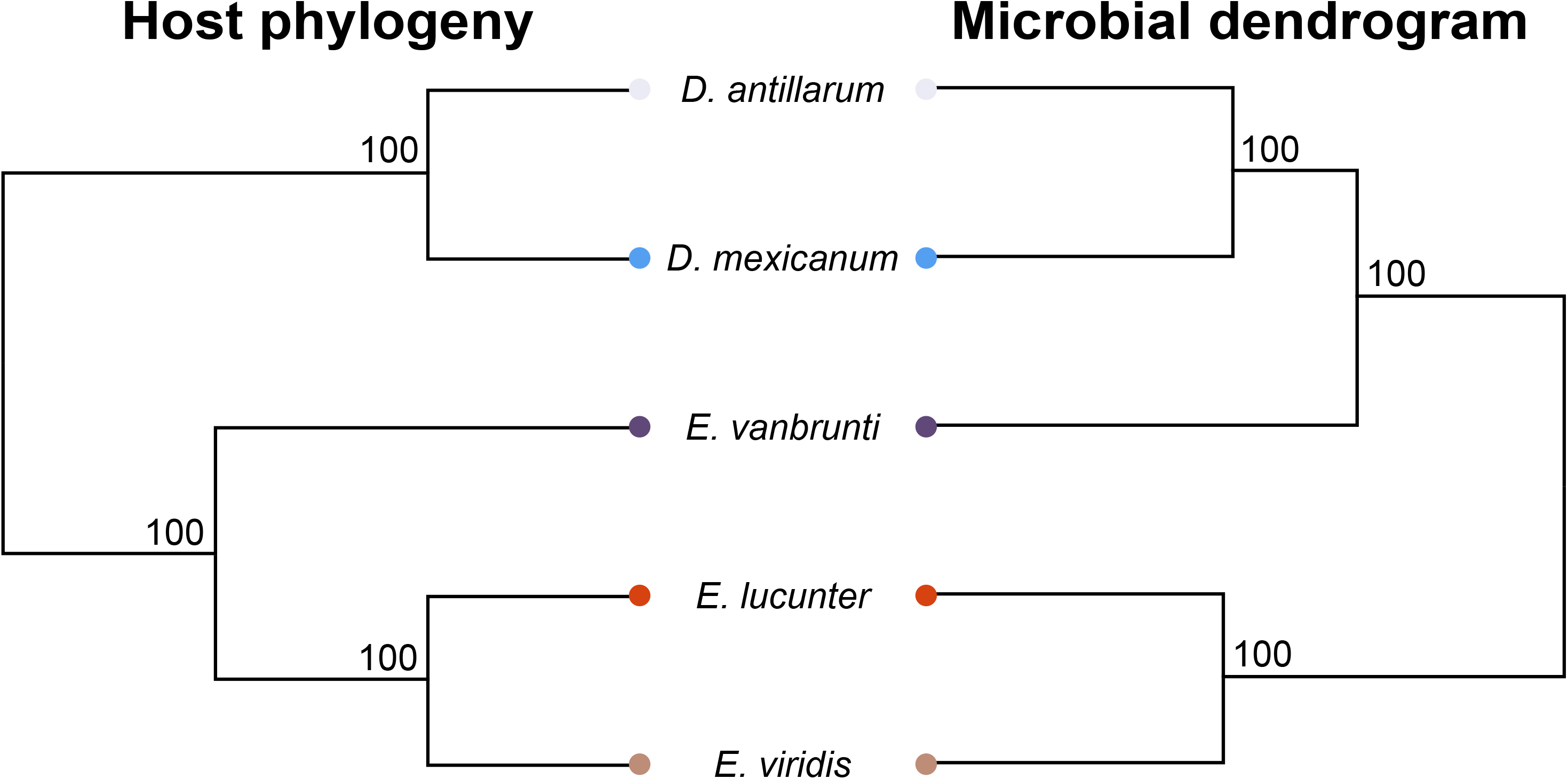
Phylosymbiosis for geminate species pairs. Gene tree using CO1 for *Echinometra* and *Diadema* geminate species pairs and microbial dendrograms based on community composition (*i.e.*, weighted UniFrac). CO1 sequences for *Echinometra* are from McCartney et al. (2000) and those for *Diadema* are from Lessios et al. (2001).

*Echinometra*-associated bacterial communities were, on average, more diverse in individual taxa and phylogenetic breadth than *Diadema* (Fig. 4A-B; Table S5). Diversity within the eggs of geminate species pairs differed between genera: those of Caribbean *Echinometra* were similar to each other but more diverse than the Pacific counterpart whereas those of the Pacific *D. mexicanum* were more diverse than the Caribbean *D. antillarum* (Fig. 4; Table S5). Moreover, the bacterial communities of *Echinometra* were more taxonomically dominant and, thus, less even than *Diadema* (Fig. 4C-D; Table S5). The eggs of all three *Echinometra* species were associated with similarly dominant bacterial communities, while the eggs of *D. mexicanum* had a more taxonomically dominant community than that of *D. antillarum* (Fig. 4; Table S5).

**Fig. 4:**
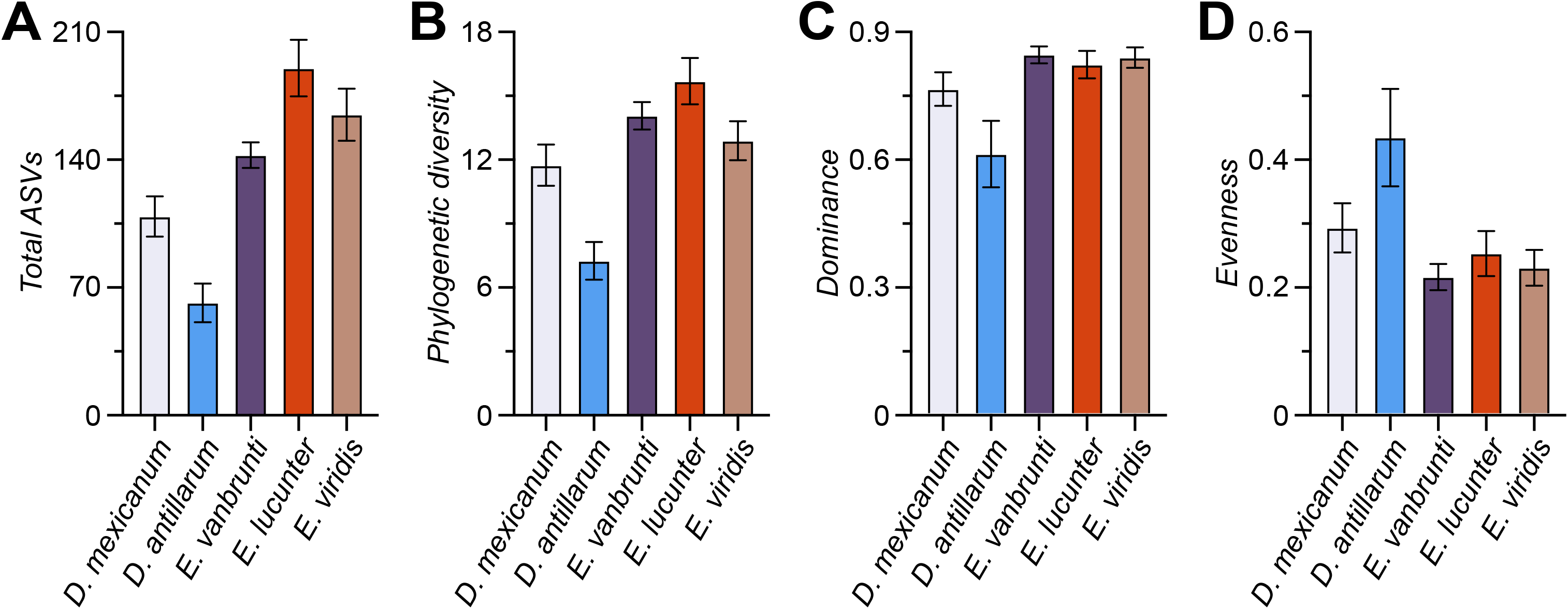
Bacterial community diversity of each sea urchin host. Diversity of the bacterial communities associated with eggs of *Echinometra* and *Diadema*, as estimated by (A) enumerating total Actual Sequence Variants, (B) Faith’s phylogenetic diversity, (C) McIntosh dominance, and (D) McIntosh evenness.

### Taxonomic representation and divergence

Microbiota of both *Echinometra* and *Diadema* eggs were primarily composed of two bacterial classes, the Bacteroidia (Bacteroidetes) and Gamaproteobacteria (Proteobacteria) (Fig. 5A). Bacteroidia, on average, represented ~24.2% and ~31.1% of the bacterial community of *Echinometra* and *Diadema*, respectively, while the Gammaproteobacteria represented ~31.9% and ~37.9% (Fig. 5A). The eggs of the Caribbean *Echinometra*, in particular, were also dominated by the Oxyphotobacteria (Phylum: Cyanobacteria), which represented ~24.8% of the community (Fig. 5A). This bacterial group was significantly more abundant in the Caribbean *Echinometra* than the Pacific *E. vanbrunti* and represented ~2-3-times more of the community (Fig 5B; Table S6). In addition to these groups, the eggs of *Diadema* had 8 other bacterial classes that represented between ~1% and ~7.6% of the community, while *Echinometra* had 7 other bacterial classes that ranged from ~1% to ~5.2% of the community (Fig 5A).

**Fig. 5:**
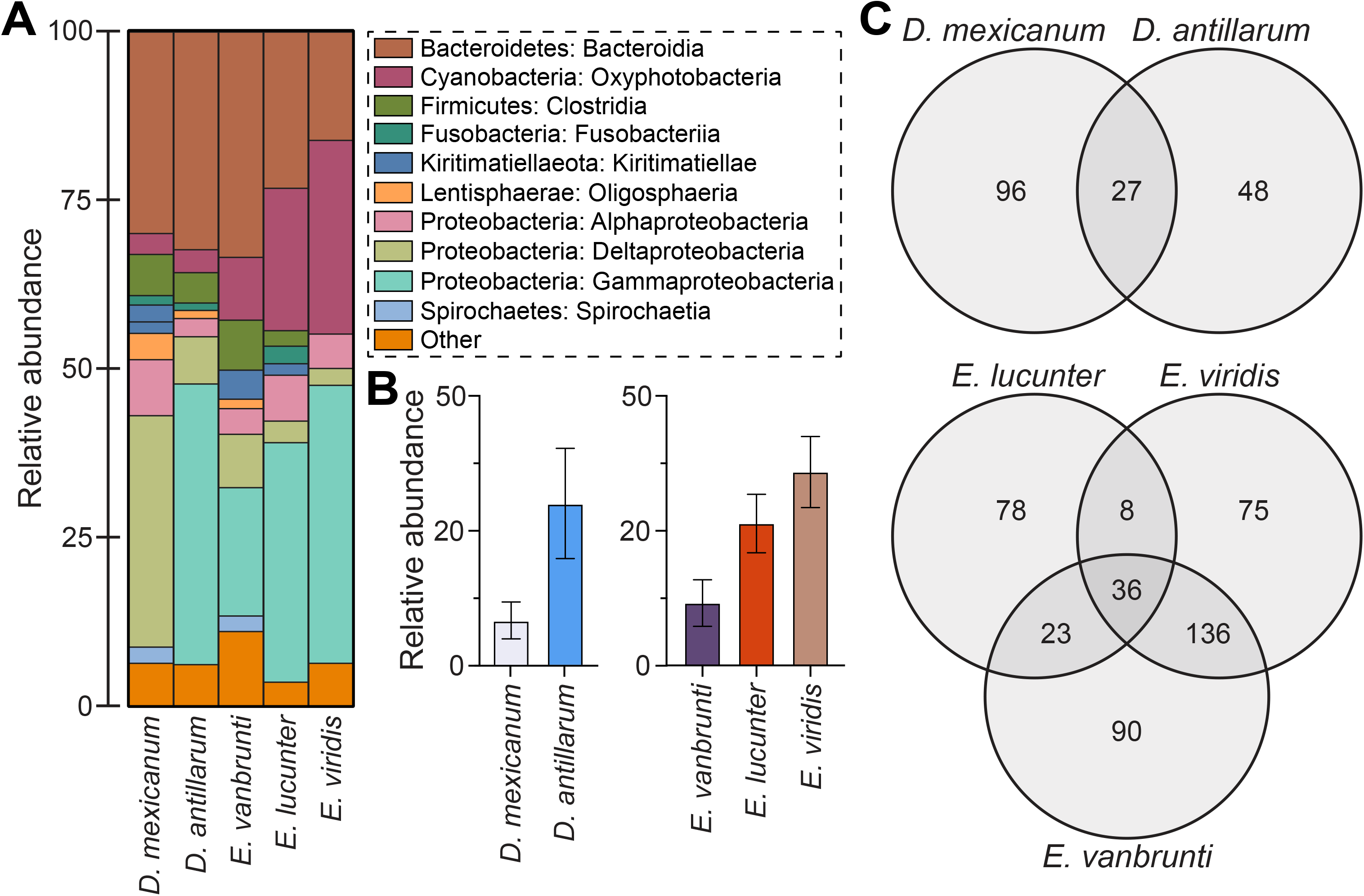
Bacterial taxa of each sea urchin species. (A) Class-level profiles of the bacterial communities associated with the *Echinometra* and *Diadema* species. (B) Differential abundance in a *Kistimonas* ASV associated with *Diadema* species (left) and Oxyphotobacteria associated with *Echinometra* species (right). (C) Venn diagrams of ASVs of the *Diadema* (top) and *Echinometra* (bottom) geminate species pairs.

There were 446 ASVs between the *Echinometra* geminate species. Of these, 90 (20.2%), 78 (17.5%), and 75 (16.8%) were specific to *E. vanbrunti*, *E. lucunter*, and *E. viridis*, respectively, while 36 (8.1%) were shared between these species (Fig. 5C, S2). Notably, a total of 136 (38.6%) ASVs were shared between the Caribbean *E. lucunter* and *E. viridis,* while 41 (9.2%) or 59 (13.2%) ASVs were shared between either species of Caribbean *Echinometra* and *E. vanbrunti* (Fig. 5C, S2). The *Diadema* geminate species, on the other hand, associated with 171 ASVs; 96 and 48 were specific to *D. mexicanum* and *D. antillarum*, respectively, and 27 were shared. *D. antillarum* also associated with the most abundant ASV, which belonged to *Kistimonas*, represented ~29.9% of the community, and was significantly more abundant in *D. antillarum* than *D. mexicanum* (Fig 5B; Table S6).

## DISCUSSION

The microbiota with which animals associate is often attributed to their evolutionary history or the environment (Carrier & Reitzel 2017, Lim & Bordenstein 2020). Studies assessing the relative importance of host’s evolutionary history or the environment suggest that both influence community composition, but one factor is commonly more pronounced. One “natural experiment” to test whether either factor or the interaction between these factors primarily drives community composition is to compare geminate species pairs (Jordan 1908). One example are echinoids separated by the Isthmus of Panama (Lessios 2008, O’Dea et al. 2016, Wilkins et al. 2019).

By comparing the egg-associated microbiota for the *Echinometra* and *Diadema* geminate species pairs, we reach three main findings. First, both pairs of geminate species associated with distinct bacterial communities that reflect a relationship consistent with phylosymbiosis. Second, the relatedness of these microbiota—based on both membership and composition—supports the hypothesis that the interaction between the animal’s evolutionary history and the environment best explains differences in these communities. Third, particular microbial taxa (*e.g.*, Oxyphotobacteria and *Kistimonas*) differed considerably between, but not within, oceans.

Like the developmental stages of many marine invertebrates, echinoid embryos and larvae associate with species-specific bacterial communities that are composed of hundreds of taxa (Carrier & Reitzel 2018, Carrier & Reitzel 2019a, b, Carrier & Reitzel 2020). Presently, no study has compared the microbiome of echinoids across their evolutionary history, but the microbiota associated with three confamilial echinoids showed a phylogenetic signal (Carrier & Reitzel 2018, Carrier & Reitzel 2019a). Multiple studies do, however, suggest that feeding environment and geographic locations with distinct oceanographic conditions influence the composition of microbiota with which echinoid larvae associate (Carrier & Reitzel 2018, Carrier et al. 2019, Carrier & Reitzel 2019b, Carrier & Reitzel 2020).

Since their separation ~2.8 million years ago, echinoid geminate species pairs have diverged in several aspects of their biology and ecology, including egg size and biochemical composition (Lessios 1990, McAlister & Moran 2012), larval feeding ecology (McAlister 2008), and reproductive ecology (Lessios 1981, 1984). In this study, we show that the bacterial communities with which they associate are another biological difference between these species pairs. Specifically, Pacific and Caribbean geminate pairs share between ~10% and ~15% of their microbiota. This fraction of shared bacterial taxa is consistent with differences between the microbiota of *Strongylocentrotus droebachiensis* larvae from multiple geographical locations (Carrier et al. 2019). Differences in these communities may reflect common taxonomic differences amongst echinoid species and/or between populations.

One exception to interspecific differences in egg microbiota is our finding regarding the two Caribbean *Echinometra* species, *E. lucunter* and *E. viridis*. These species diverged from each other less than ~1.6 million years ago (McCartney et al. 2000); after this time interval they still maintain a similarity of ~55% of their bacterial taxa. This lower level of taxonomic divergence has not resulted in a specific-specific bacterial community, which is hypothesized to be a fundamental property of animal-associated microbiota (Gilbert et al. 2012, McFall-Ngai et al. 2013, Bordenstein & Theis 2015). Recently, however, notions for universal rules governing animal-microbe symbioses have been called into question (Hammer et al. 2019). The comparison between these two Caribbean *Echinometra* species may provide initial evidence against the universality of species-specific microbiota.

In marine and terrestrial taxa, the evolutionary history of the host tends to mirror the relatedness of the microbial community (*i.e.*, phylosymbiosis; Brooks et al. 2016, Lim & Bordenstein 2020). When comparing host phylogeny and microbial dendrograms for the *Echinometra* and *Diadema* geminate species pairs, we find evidence for phylosymbiosis (Brooks et al. 2016, Lim & Bordenstein 2020). Although these trees were not fully congruent, the evolutionary signal was strong enough to be detected, despite the environmental influence in this system. Relatedness of these bacterial communities did not fully reflect either factor; instead, what was observed was an intermediate between a host- and environment-driven pattern.

Symbioses between animals and microbes are a product of the interaction between the host genotype (G_H_), the microbial metagenome (G_M_), and the environment (E) (Zilber-Rosenberg & Rosenberg 2008, Bordenstein & Theis 2015, Carrier & Reitzel 2017). This tripartite interaction (G_H_ × G_M_ × E) was evident in the *Echinometra* and *Diadema* geminate species pairs, where the Pacific *E. vanbrunti* grouped with *Diadema* geminate species pair and the two Caribbean *Echinometra* species grouped separately. Provided that both evolutionary history and environment contribute to the composition of echinoid bacteria, the known history of the geminate species can shed light on an unresolved question of bacterial symbiosis: which symbionts have a deeper common history with the host (*e.g.*, co-speciation; Peek et al. 1998, Funkhouser & Bordenstein 2013, Moeller et al. 2016), and which partnerships formed as the physical and biological properties of these ocean drastically changed (Lessios 2008, O’Dea et al. 2016)?

Two potential candidates that may be traced with host evolution or be environmentally relevant are the Oxyphotobacteria and *Kistimonas*. Oxyphotobacteria are a group of cyanobacteria that perform oxygenic photosynthesis (Soo et al. 2017). In our data, this bacterial class was ~2-3-times as abundant in the Caribbean *Echinometra.* The Caribbean is oligotrophic relative to the eastern Pacific, which has been hypothesized to drive the evolution of a number of life history traits, presumed to be adaptations to its lower productivity (Lessios 2008). Multiple echinoderms living in oligotrophic seas have been observed to associate with bacterial lineages known to perform photosynthesis (Bosch 1992, Galac et al. 2016, Carrier et al. 2018, Carrier & Reitzel 2020); however, the function of these bacteria remain unknown. *Kistimonas* is a recently identified bacterial lineage known to associate with various marine invertebrates (Choi et al. 2010, Lee et al. 2012, Ellis et al. 2019). The symbiotic functions of the *Kistimonas* remains unknown, but the dominance of this bacterium on the eggs of *D. antillarum*, but not *D. mexicanum*, may suggest some means to cope with an oligotrophic sea.

Taken together, data presented here support a hypothesis that the bacterial communities of echinoid geminate species pairs have diverged considerably since the formation of the Isthmus of Panama and may be a product of both the evolutionary history of the host and subsequent evolution in their respective environments. The functional importance of these bacterial communities and, consequently, whether they are adapted for each oceanographic regime remains an open question (Wilkins et al. 2019). Functional potential and expression profiles of these microbial communities may be assessed using meta-genomics and meta-transcriptomics, which may then be coupled with culturing of individual bacterial taxa and performing add-back experiments to test whether performance is enhanced under different environments (Moitinho-Silva et al. 2014, Slaby et al. 2017, Domin et al. 2018, Carrier & Reitzel 2020).

## Acknowledgements

We thank the staff of the Smithsonian Tropical Research Institute and Axel and Ligia Calderon and Laura Geyer for their valuable assistance with experimental logistics; Karen Lopez and Daniel Janies (UNC Charlotte) for sequencing resources and technical assistance with sequencing. T.J.C. was supported by an NSF Graduate Research Fellowship and a Smithsonian Institute graduate student fellowship; H.L. was supported by General Research Funds from STRI; and A.M.R. was supported by a Human Frontier Science Program Award (RGY0079/2016).

---

**Fig. S1:**
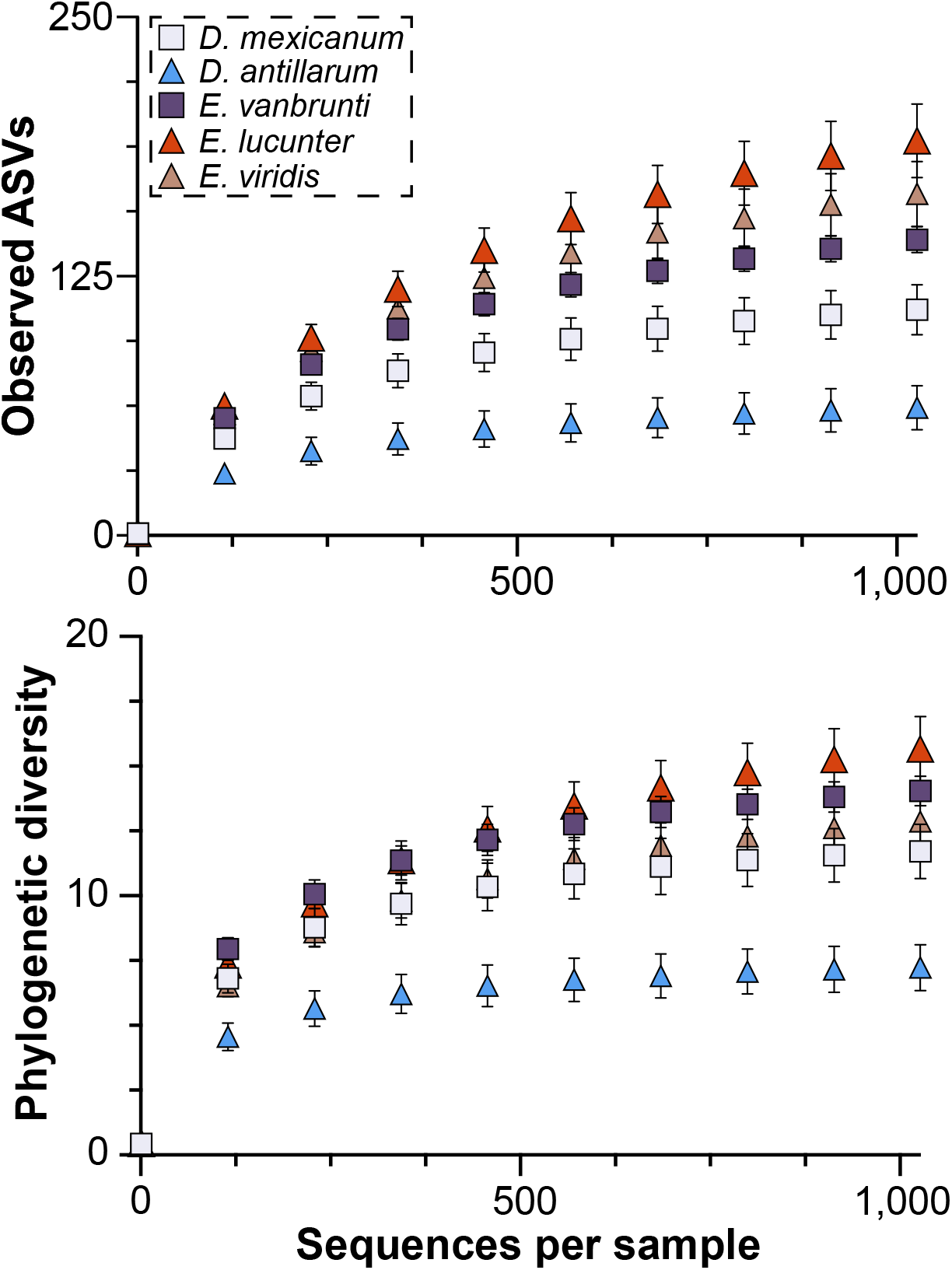
Alpha rarefaction curve for *Diadema* and *Echinometra* species. Alpha rarefaction curve for the bacterial community associated with *D. mexicanum*, *D. antillarum*, *E. vanbrunti*, *E. lucunter*, and *E. viridis* eggs. This was based on a rarefaction depth of 1,027 sequences and was used for all analyses.

**Fig. S2:**
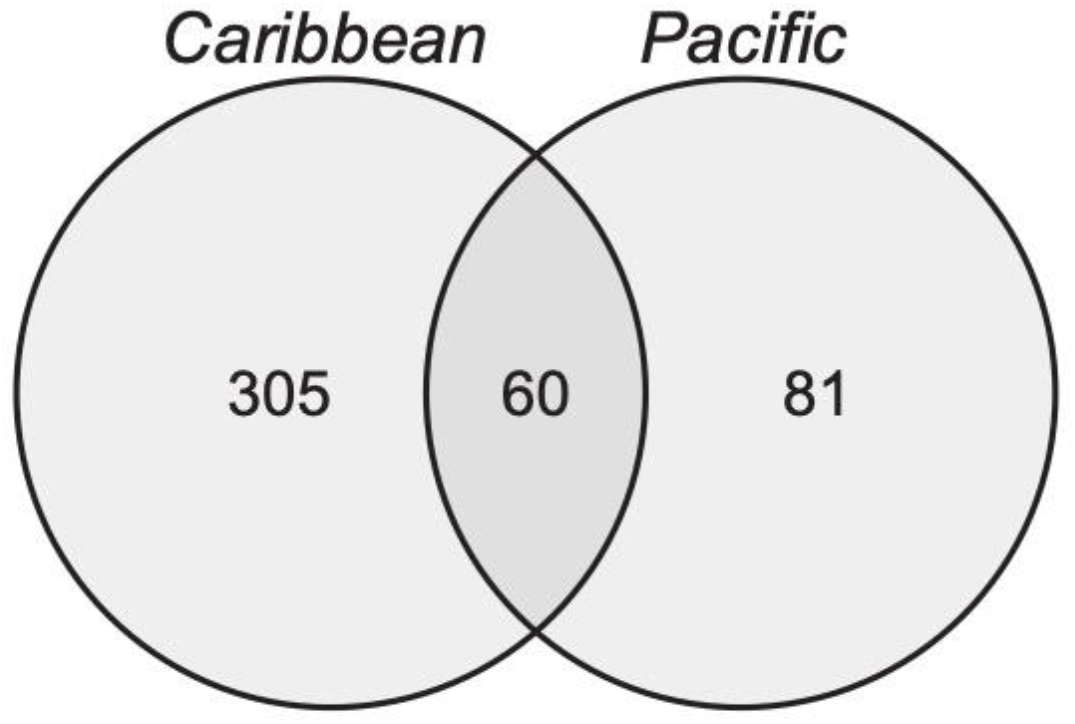
Taxonomic differences between the egg microbiota of *Echinometra* geminate species. Differences in the taxonomic representation for *Echinometra* species from the Caribbean (*E. lucunter* and *E. viridis* together) and the Pacific (*E. vanbrunti*).

## References

Aldred N, Nelson A (2019) Microbiome acquisition during larval settlement of the barnacle Semibalanus balanoides. Biology Letters 15:20180763

Amir A, McDonald D, Navas-Molina JA, Kopylova E, Morton JT, Xu ZZ, Kightley EP, Thompson LR, Hyde ER, Gonzalez A, Knight R (2017) Deblur rapidly resolves single-nucleotide community sequence patterns. mSystems 2:e00191–00116

Bogdanowicz D, Giaro K, Wróbel B (2012) TreeCmp: comparison of trees in polynomial time. Evolutionary Bioinformatics 8:475–487

Bolyen E, Rideout JR, Dillon MR, Bokulich NA, Abnet C, Al-Ghalith GA, Alexander H, Alm EJ, Arumugam M, Asnicar F, Bai Y, Bisanz JE, Bittinger K, Brejnrod A, Brislawn CJ, Brown CT, Callahan BJ, Caraballo-Rodríguez AM, Chase J, Cope E, Da S.ilva R, Dorrestein PC, Douglas GM, Durall DM, Duvallet C, Edwardson CF, Ernst M, Estaki M, Fouquier J, Gauglitz JM, Gibson DL, Gonzalez A, Gorlick K, Guo J, Hillmann B, Holmes S, Holste H, Huttenhower C, Huttley G, Janssen S, Jarmusch AK, Jiang L, Kaehler B, Kang KB, Keefe CR, Keim P, Kelley ST, Knights D, Koester I, Kosciolek T, Kreps J, Langille MG, Lee J, Ley R, Liu Y, Loftfield E, Lozupone C, Maher M, Marotz C, Martin BD, McDonald D, McIver LJ, Melnik AV, Metcalf JL, Morgan SC, Morton J, Naimey AT, Navas-Molina JA, Nothias LF, Orchanian SB, Pearson T, Peoples SL, Petras D, Preuss ML, Pruesse E, Rasmussen LB, Rivers A, Robeson, II MS, Rosenthal P, Segata N, Shaffer M, Shiffer A, Sinha R, Song SJ, Spear JR, Swafford AD, Thompson LR, Torres PJ, Trinh P, Tripathi A, Turnbaugh PJ, Ul-Hasan S, van d.er H.ooft JJ, Vargas F, Vázquez-Baeza Y, Vogtmann E, von H.ippel M, Walters W, Wan Y, Wang M, Warren J, Weber KC, Williamson CH, Willis AD, Xu ZZ, Zaneveld JR, Zhang Y, Zhu Q, Knight R, Caporaso JG (2019) Reproducible, interactive, scalable and extensible microbiome data science using QIIME 2. Nat Biotechnol 37:852–857

Bordenstein SR, Theis KR (2015) Host biology in light of the microbiome: ten principles of holobionts and hologenomes. PLoS Biol 13:e1002226

Bosch I (1992) Symbiosis between bacteria and oceanic clonal sea star larvae in the western North Atlantic Ocean. Marine Biology 114:495–502

Brooks AW, Kohl KD, Brucker RM, van Opstal EJ, Bordenstein SR (2016) Phylosymbiosis: relationships and functional effects of microbial communities across host evolutionary history. PLoS Biol 14:e2000225

Callahan BJ, McMurdie PJ, Holmes SP (2017) Exact sequence variants should replace operational taxonomic units in marker-gene data analysis. The ISME Journal 11:2639–2643

Caporaso JG, Kuczynski J, Stombaugh J, Bittinger K, Bushman FD, Costello EK, Fierer N, Pena AG, Goodrich JK, Gordon JI, Huttley GA, Kelley ST, Knights D, Koenig JE, Ley RE, Lozupone CA, McDonald D, Muegge BD, Pirrung M, Reeder J, Sevinsky JR, Turnbaugh PJ, Walters WA, Widmann J, Yatsunenko T, Zaneveld J, Knight R (2010) QIIME allows analysis of high-throughput community sequencing data. Nat Methods 7:335–336

Carrier TJ, Dupont S, Reitzel AM (2019) Geographic location and food availability offer differing levels of influence on the bacterial communities associated with larval sea urchins. FEMS Microbiol Ecol 95:fiz103

Carrier TJ, Reitzel AM (2017) The hologenome across environments and the implications of a host-associated microbial repertoire. Frontiers in Microbiology 8:802

Carrier TJ, Reitzel AM (2018) Convergent shifts in host-associated microbial communities across environmentally elicited phenotypes. Nature Communications 9:952

Carrier TJ, Reitzel AM (2019a) Bacterial community dynamics during embryonic and larval development of three confamilial echinoids. Mar Ecol Prog Ser 611:179–188

Carrier TJ, Reitzel AM (2019b) Shift in bacterial taxa precedes morphological plasticity in a larval echinoid. Marine Biology 166:164

Carrier TJ, Reitzel AM (2020) Symbiotic life of echinoderm larvae. Frontiers in Ecology and Evolution 7:509

Carrier TJ, Wolfe K, Lopez K, Gall M, Janies DA, Byrne M, A.M. R (2018) Diet-induced shifts in the crown-of-thorns (*Acanthaster* sp.) larval microbiome. Marine Biology 165:157

Choi EJ, Kwon HC, Sohn YC, Yang HO (2010) *Kistimonas asteriae* gen. nov., sp. nov., a gammaproteobacterium isolated from *Asterias amurensis*. International Journal of Systematic and Evolutionary Microbiology 60:938–943

Domin H, Zurita-Gutiérrez YH, Scotti M, Buttlar J, Hentschel U, Fraune S (2018) Predicted bacterial interactions affect in vivo microbial colonization dynamics in *Nematostella*. Frontiers in Microbiology 9:728

Drummond AJ, Suchard MA, Xie D, Rambaut A (2012) Bayesian Phylogenetics with BEAUti and the BEAST 1.7. Mol Biol Evol 29:1969–1973

Easson CG, Thacker RW (2014) Phylogenetic signal in the community structure of host-specific microbiomes of tropical marine sponges. Frontiers in Microbiology 5:532

Ellis JC, Thomas MS, Lawson PA, Patel NB, Faircloth W, Hayes SE, Linton EE, Norden DM, Severenchuk IS, West CH, Brown JW, Plante RG, Plante CJ (2019) *Kistimonas alittae* sp. nov., a gammaproteobacterium isolated from the marine annelid *Alitta succinea*. International Journal of Systematic and Evolutionary Microbiology 69:235–240

Funkhouser L, Bordenstein SR (2013) Mom knows best: the universitality of maternal microbial transmission. PLoS Biol 11:e1001631

Galac MR, Bosch I, Janies DA (2016) Bacterial communities of oceanic sea star (Asteroidea: Echinodermata) larvae. Marine Biology 163:162

Gilbert SF, Sapp J, Tauber AI (2012) A symbiotic view of life: we have never been individuals. Q Rev Biol 87:325–341

Hammer TJ, Sanders JG, Fierer N (2019) Not all animals need a microbiome. FEMS Microbiol Lett 336:fnz117

Hickerson MJ, Stahl EA, Lessios HA (2006) Test for simultaneous divergence using approximate Bayesian computation. Evolution 60:2435–2453

Jordan DS (1908) The law of the geminate species. The American Naturalist 42:73–80

Klindworth A, Pruesse E, Schweer T, Peplies J, Quast C, Horn M, Glockner FO (2013) Evaluation of general 16S ribosomal RNA gene PCR primers for classical and next-generation sequencing-based diversity studies. Nucleic Acids Research 41:e1

Kohl KD, Carey HV (2016) A place for host-microbe symbiosis in the comparative physiologist’s toolbox. Journal of Experimental Biology 219:3496–3504

Lee J, Shin N-R, Lee H-W, Roh SW, Kim M-S, Kim Y-O, Bae J-W (2012) *Kistimonas scapharcae* sp. nov., isolated from a dead ark clam (*Scapharca broughtonii*), and emended description of the genus *Kistimonas*. International Journal of Systematic and Evolutionary Microbiology 62:2865–2869

Lessios HA (1981) Reproductive periodicity of the echinoids *Diadema* and *Echinometra* on the two coasts of Panama. Journal of Experimental Marine Biology and Ecology 50:47–61

Lessios HA (1984) Possible prezygotic reproductive isolation in sea urchins separated by the Isthmus of Panama. Evolution 38:1144–1148

Lessios HA (1990) Adaptation and phylogeny as determinants of egg size in echinoderms from the two sides of the Isthmus of Panama. The American Naturalist 135:1–13

Lessios HA (2008) The great american schism: divergence of marine organisms after the rise of the Central American Isthmus. Annual Review of Ecology, Evolution, and Systematics 39:63–91

Lessios HA, Kessing BD, Pearce JS (2001) Population structure and speciation in tropical seas: global phylogeography of the sea urchin *Diadema*. Evolution 55:955–975

Lim SJ, Bordenstein SR (2020) An introduction to phylosymbiosis. Proceedings of the Royal Society B 287:20192900

Lozupone C, Knight R (2005) UniFrac: a new phylogenetic method for comparing microbial communities. Applied and Environmental Microbiology 71:8228–8235

McAlister JS (2008) Evolutionary responses to environmental heterogeneity in Central American echinoid larvae: plastic versus constant phenotypes. Evolution 62:1358–1372

McAlister JS, Moran AL (2012) Relationships among egg size, composition, and energy: a comparative study of geminate sea urchins. PLoS ONE 7:e41599

McCartney MA, Kelley G, Lessios HA (2000) Dispersal barriers in tropical oceans and speciation in Atlantic and eastern Pacific sea urchins of the genus *Echinometra*. Molecular Ecology 9:1391–1400

McFall-Ngai M, Hadfield MG, Bosch TCG, Carey HV, Domazet-Loso T, Douglas AE, Dubilier N, Eberl G, Fukami T, Gilbert SF, Hentschel U, King N, Kjelleberg S, Knoll AH, Kremer N, Mazmanian SK, Metcalf JL, Nealson K, Pierce NE, Rawls JF, Reid A, Ruby EG, Rumpho M, Sanders JG, Tautz D, Wernegreen JJ (2013) Animals in a bacterial world, a new imperative for the life sciences. P Natl Acad Sci USA 110:3229–3236

Moeller AH, Caro-Quintero A, Mjungu D, Georgiev AV, Lonsdorf EV, Muller MN, Pusey AE, Peeters M, Hahn BH, Ochman H (2016) Cospeciation of gut microbiota with hominids. Science 353:380–382

Moitinho-Silva L, Seridi L, Ryu T, Voolstra CR, Ravasi T, Hentschel U (2014) Revealing microbial functional activities in the Red Sea sponge *Stylissa carteri* by metatranscriptomics. Environmental Microbiology 16:3683–3698

Mortzfeld BM, Urbanski S, Reitzel AM, Kunzel S, Technau U, Fraune S (2015) Response of bacterial colonization in *Nematostella vectensis* to development, environment and biogeography. Environmental Microbiology 18:1764–1781

O’Brien PA, Webster NS, Miller DJ, Bourne DG (2019) Host-microbe coevolution: applying evidence from model systems to complex marine invertebrate holobionts. mBio 10:e02241–02218

O’Dea A, Lessios HA, Coates AG, Eytan RI, Restrepo-Moreno SA, Cione AL, Collins LS, de Queiroz A, D.W. F, Norris RD, Stallard RF, Woodburne MO, Aguilera O, Aubry M-P, Berggren WA, Budd AF, Cozzuol MA, Coppard SE, Duque-Caro H, Finnegan S, Gasparini GM, Grossman EL, Johnson KG, Keigwin LD, Knowlton N, Leigh EG, Leonard-Pingel JS, Marko PB, Pyenson ND, Rachello-Dolmen PG, Soibelzon E, Soibelzon L, Todd JA, Vermeij GJ, Jackson JBC (2016) Formation of the Isthmus of Panama. Sci Adv 2:e1600883

Peek AS, Feldman RA, Lutz RA, Vrijenhoek RC (1998) Cospeciation of chemoautotrophic bacteria and deep sea clams. P Natl Acad Sci USA 95:9962–9966

Pollock FJ, McMinds R, Smith S, Bourne DG, Willis BL, Medina M, Vega Thurber R, Zaneveld JR (2018) Coral-associated bacteria demonstrate phylosymbiosis and cophylogeny. Nature Communications 9:4912

Quast C, Pruesse E, Yilmaz P, Gerken J, Schweer T, Yarza P, Peplies J, Glockner FO (2013) The SILVA ribosomal RNA gene database project: improved data processing and web-based tools. Nucleic Acids Research 41:590–596

Rambaut A, Drummond AJ, Xie D, Baele G, Suchard MA (2018) Posterior summarization in Bayesian phylogenetics using tracer 1.7. Systematic Biology 67:901–904

Rognes T, Flouri T, Nichols B, Quince C, Mahé F (2016) VSEARCH: a versatile open source tool for metagenomics. PeerJ 4:e2584

Schmitt S, Tsai P, Bell J, Fromont J, Ilan M, Lindquist N, Perez T, Rodrigo A, Schupp PJ, Vacelet J, Webster N, Hentschel U, Taylor MW (2012) Assessing the complex sponge microbiota: core, variable and species-specific bacterial communities in marine sponges. ISME Journal 6:564–576

Slaby BM, Hackl T, Horn H, Bayer K, Hentschel U (2017) Metagenomic binning of a marine sponge microbiome reveals unity in defense but metabolic specialization. The ISME Journal 11:2465–2478

Soo RM, Hemp J, Parks DH, Fischer WW, Hugenholtz P (2017) On the origins of oxygenic photosynthesis and aerobic respiration in Cyanobacteria. Science 355:1436–1440

Soto W, Gutierrez J, Remmenga MD, Nishiguchi MK (2009) Salinity and temperature effects on physiological responses of *Vibrio fischeri* from diverse ecological niches. Microb Ecol 57:140–150

Thomas T, Moitinho-Silva L, Lurgi M, Bjork JR, Easson C, Astudillo-García C, Olson JB, Erwin PM, López-Legentil S, Luter H, Chaves-Fonnegra A, Costa R, Schupp PJ, Steindler L, Erpenbeck D, Gilbert J, Knight R, Ackermann G, Lopez JV, Taylor MW, Thacker RW, Montoya JM, Hentschel U, Webster NS (2016) Diversity, structure and convergent evolution of the global sponge microbiome. Nature Communications 7:11870

Webster NS, Botte ES, Soo RM, Whalan S (2011) The larval sponge holobiont exhibits high thermal tolerance. Environmental Microbiology Reports 3:756–762

Wilkins LGE, Leray M, O’Dea A, Yuen B, Peixoto RS, Pereira TJ, Bik HM, Coil DA, Duffy JE, Herre EA, Lessios HA, Lucey NM, Mejia JC, Rasher DB, Sharp KH, Sogin EM, Thacker RW, Thurber RV, Wcislo WT, Wilbanks EG, Eisen JA (2019) Host-associated microbiomes drive structure and function of marine ecosystems. PLoS Biol 17:e3000533

Zilber-Rosenberg I, Rosenberg E (2008) Role of microorganisms in the evolution of animals and plants: the hologenome theory of evolution. Fems Microbiol Rev 32:723–735

